# Metabolism-weighted brain connectome reveals synaptic integration and vulnerability to neurodegeneration

**DOI:** 10.1101/2025.10.02.679973

**Authors:** Mahnaz Ashrafi, Laura Fraticelli, Gabriel Castrillón, Valentin Riedl

## Abstract

The brain’s capacity for integration arises from both its structural wiring and energetically demanding electrochemical signaling. Yet current connectome analyses treat network nodes as functionally homogeneous, ignoring that neural communication is constrained by metabolic cost. Here we introduce a metabolism-weighted connectome, a fully weighted brain graph in which both connections and the metabolic activity of each node describe the network’s capacity for integration. Using three datasets of simultaneous fMRI and FDG-PET acquisitions, we define metabolism-weighted centrality (MwC), a biologically grounded index of each region’s signaling dominance that integrates functional connectivity (FC) with local energy metabolism. MwC provides a more accurate representation of cortical activity flow than classical edge-based metrics and reveals that metabolically active hubs align with higher-order cognitive networks. Transcriptomic and synaptic imaging data demonstrate that these hubs exhibit increased synaptic energy turnover, linking activity-driven centrality to the molecular architecture of signaling. Notably, the same high-MwC regions show greater susceptibility to neurodegenerative pathology, suggesting that lifelong metabolic demand influences both integrative function and disease vulnerability. By linking neuronal metabolism to network organization, our framework bridges cellular energetics and system-level computation, opening new avenues for interpreting brain vulnerability and performance.

**Significance Statement:** The various regions of the brain exhibit unequal energy consumption. Specifically, areas associated with cognitive functions such as memory and attention are metabolic hotspots of intense neural activity. Yet standard connectivity analyses ignore this energetic dimension, treating all regions as functionally equivalent. We introduce the metabolism-weighted connectome (MwC), a framework that maps brain connectivity weighted by each region’s energy expenditure. Using simultaneous PET-MRI in healthy volunteers, we show that energetically most active hubs anchor networks for complex cognition, exhibit molecular signatures of intense synaptic activity, and are disproportionately vulnerable to Alzheimer’s disease. These findings establish a unifying principle: the brain regions most essential for cognition bear the greatest metabolic burden, and this cost may underlie their susceptibility to neurodegeneration.

## INTRODUCTION

The brain’s ability to sustain complex cognition depends on the coordinated activity of distributed networks. These networks, classically represented as connectomes, describe how anatomically distinct regions interact through structural and functional links (1, 2). While network science has transformed our understanding of brain organization (3–5), most existing models describe how regions are connected, not how intensively they communicate. The energetic cost of maintaining these interactions, a defining constraint of neural computation, remains largely absent from current connectome frameworks.

Conventional graph representations, which assign weights to edges, evaluate a region’s importance by its pairwise connectivity strength or brain-wide strength centrality (6–8). This edge-focused perspective implicitly assumes that all nodes operate with similar activity and metabolic demand. Yet, neural signaling is profoundly heterogeneous across the cortex, with metabolic activity varying regionally and in brain disorders (9–13). As a result, regions defined as “hubs” by topological criteria may, in biological terms, contribute little to actual signaling, whereas metabolically dominant but sparsely connected regions may play critical integrative roles in the connectome. This conceptual gap limits our ability to relate network architecture to the physiological constraints of neuronal function. Recent developments in network theory have sought to address such limitations by incorporating node attributes, such as activity or functional diversity, into graph formulations (14–19). Extending this logic to the brain implies that an accurate connectome must incorporate both connection strength and local signaling activity.

The brain relies on a continuous supply of glucose as its primary energy substrate. Neuronal energy expenditure is dominated by signaling activity, which accounts for an estimated 75% of grey matter energy consumption, with postsynaptic potentials alone representing half of that fraction (20–22). Because signaling is among the most energetically demanding processes in the brain, regional metabolic rate serves as a direct and quantitative proxy for the intensity of local neuronal communication. [¹⁸F]Fluorodeoxyglucose positron emission tomography (FDG-PET) allows to measure the cerebral metabolic rate of glucose (CMRglc) in vivo, providing a regionally resolved index of signaling activity (23, 24). FDG-PET uptake thus reflects the energetic load at individual nodes of the brain connectome and the extent to which a region engages in information exchange. Functional connectivity (FC), by contrast, captures the temporal coherence of blood-oxygen-level-dependent (BOLD) signals between distributed cortical regions, indexing the statistical coupling of moment-to-moment activity between node pairs (25, 26). While FC identifies which regions communicate at the time of measurement, it remains blind to the absolute intensity of that exchange: BOLD amplitude changes are inherently relative, and FC edge weights reflect statistical dependence rather than the extent of communication. Whether regions of high network centrality also incur a proportionally higher metabolic cost is therefore an open question.

Here we introduce the metabolism-weighted connectome, a biologically informed network model that integrates connectivity and regional energy metabolism to capture the brain’s activity-constrained architecture. Within this framework, we define metabolism-weighted centrality (MwC) as a quantitative measure of signaling dominance reflecting a region’s connectivity weighted by the metabolic activity of its partners. Concretely, a region attains high MwC not merely by being densely connected, but by being preferentially coupled to regions that themselves sustain high rates of glucose metabolism. Using three datasets with simultaneous FDG-PET and fMRI acquisitions, we show that MwC identifies metabolically dominant hubs closely linked to cognitive networks, synaptic density, and neurodegenerative vulnerability. This model provides a mechanistic principle for understanding how energetic costs shape network integration, revealing that the very regions enabling high cognitive complexity also bear the metabolic burden that predisposes them to disease.

## RESULTS

### Metabolism-weighted centrality (MwC) defines signaling dominance

To move beyond a topology-based description of brain networks, we sought to incorporate energetic information into the connectome. Using simultaneously acquired FDG-PET and fMRI data, we developed a fully-weighted brain graph, in which both nodes and edges have metabolic and functional weights. We analyzed three independent datasets, each acquired on an integrated PET/MRI scanner at two different sites (TUM University Hospital Munich, Medical University of Vienna). The main dataset included 20 healthy subjects (10 females, age: 34.5 ± 5 years, TUM), and the two replication datasets comprised 10 subjects each (5 females, age: 27 ± 5 years, TUM.rep; 5 females, age: 27 ± 7 years, Vienna). All datasets provide simultaneously acquired measures of CMRglc from FDG-PET and of FC from fMRI data. We performed all analyses in subject space and modeled the brain graph on regions as defined by the Human Connectome Project Multimodal Parcellation (HCP MMP) atlas (Fig. 1a) (27).

**Fig 1:**
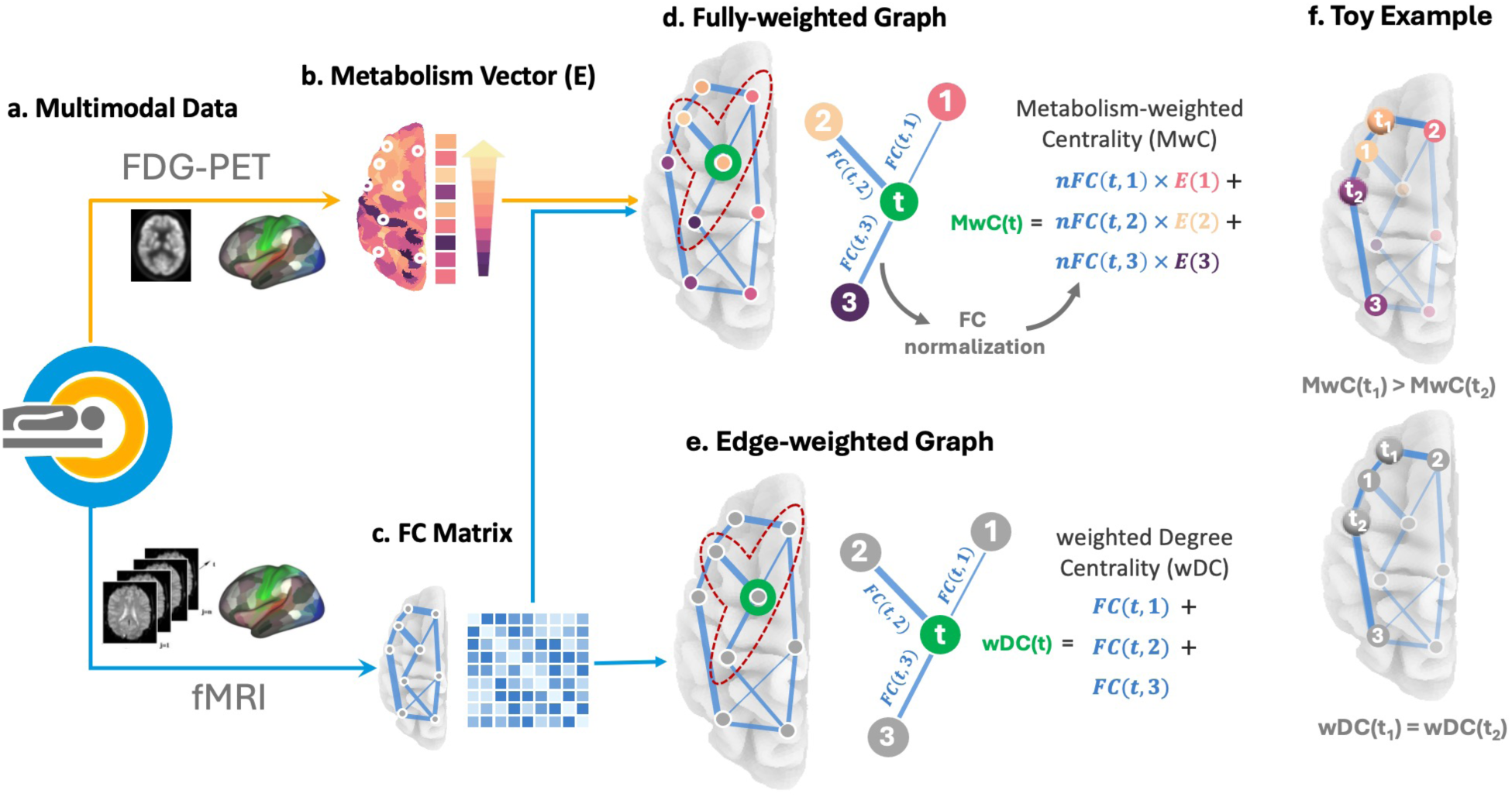
The metabolism-weighted connectome. **(a)** FDG-PET (orange) and fMRI (blue) data were acquired simultaneously on an integrated PET/MRI scanner and parcellated according to 360 ROIs from the Human Connectome Project Multi-Modal Parcellation (HCP MMP) atlas. **(b)** Brain surface representation of energy metabolism (CMRglc) across the cortex. Each white circle represents a region where the median CMRglc value is extracted and stored in a vector of energy metabolism *E*, color-coding the metabolic activity. **(c)** A network graph overlaid on the brain illustrates the brain FC profile. Nodes represent brain regions, with edge thickness indicating the weighted FC strength, calculated as the statistical dependencies between pairwise nodal fMRI time series and plotted as an FC matrix. The connectivity matrix is used to calculate both the MwC and wDC. **d)** Fully-weighted graph where the color-coded node values correspond to the values of vector *E* in b) and its edges, the FC matrix in c). The inset illustrates the MwC of a target region t in green, connected to three regions labeled 1-3, highlighted by the dashed red contour, as the sum of the product between *E(i)* and *nFC(t,i)* across its connected nodes. **(e)** Using an edge-weighted graph, the wDC is calculated as the sum of the FC of all connected nodes from fMRI data only. **(f)** Case scenario of two regions, t_1_ and t_2,_ with identical wDC values (lower panel) but different metabolic profiles. Both nodes are connected to region 1, but region t_1_ is additionally connected to 2, and region t_2_ to region 3, which have different metabolism profiles (*E(2) > E(3)*), shown in the upper panel. Integrating nodal metabolism into the model yields different MwC (MwC(t_1_) > MwC(t_2_)) while they have identical wDC (wDC(t_1_) = wDC(t_2_)).

The fully-weighted brain graph is defined as a two-layer network, including the energy metabolism in each region (Fig. 1b) and the FC matrix of correlated BOLD signal time-series for each pair of regions (Fig. 1c). We then calculated MwC for each node *t* as:

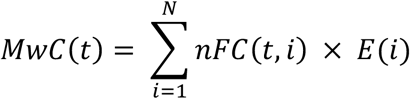

where *t* is the target region, *N* is the total number of regions connected to node *t*, *nFC(t,i)* is the normalized FC between nodes *t* and *i*, and *E(i)* is the energy metabolism of any connected region *i. nFC* is calculated by dividing the FC value between nodes *t* and *i* by the total sum of connections linked to node *t*. This normalization reflects the relative influence of each connected region *i* on the activity of region *t* (see Methods).

The region-wise calculation of MwC results from a fully-weighted brain graph that incorporates both the connectivity profile and the metabolic activity of connected regions (Fig. 1d). In contrast, the classical model of weighted degree centrality (wDC) is calculated solely as the sum of weighted edges for a given region *t*, without considering varying levels of activity in connected nodes (Fig. 1e). The toy example in Fig. 1f illustrates the critical difference between the two centrality measures for two arbitrary nodes, *t_1_* and *t_2_*. Using an edge-weighted model only, the two nodes have the same wDC and are thus ranked with equal centrality (lower panel). MwC, on the other hand, additionally considers the level of activity in connected regions 1 to 3, which results in a biologically more realistic model of information processing along the network. In our example, *t_1_* has a higher centrality within the brain connectome as it connects to nodes with higher activity than *t_2_* (upper panel).

### MwC reveals energy-based architecture of cortical integration

After calculating both graph metrics, we examined how well each centrality measure matches the signaling-related energy metabolism across the cortex. We calculated the correlations of MwC and wDC with CMRglc across brain regions and compared their performance. It is important to note that the calculation of MwC for node *t* is independent of *E(t)*, as it only uses the CMRglc from connected regions *i*. On the group level, the subject average CMRglc shows a significant positive correlation with MwC (r = 0.63, p-spin < 0.001, scatter plot in Fig. 2a) and a weaker correlation with wDC (r = 0.31, p-spin < 0.001, Fig. 2b). The group surface and graph plots in Fig. 2 illustrate the varying distribution of hub regions for MwC and wDC. While MwC hubs are located in medial and lateral parietal and lateral frontal regions, wDC hubs occur mainly in sensory-motor regions (see Supplementary Fig. S1 for additional details). The stronger association of the fully-weighted MwC compared to edge-oriented wDC in relation to CMRglc is also evident at the subject level, as shown in the violin plots of Fig. 2. MwC values range from r = 0.30 - 0.62 (all p < 0.05), while the wDC values range from r = 0.10 - 0.27 (for significant cases only). Finally, we performed identical analyses on the two independent replication datasets and observed similarly strong (r = 0.51 – 0.57) correlations for MwC with CMRglc (see Supplementary Fig. S2). In addition to wDC, we also tested another prominent edge-oriented centrality measure, participation coefficient (PC), to explain signaling-related energy metabolism. Similarly to wDC, PC showed a weaker correlation with CMRglc (r = 0.29, p-spin < 0.001) than MwC (see Supplementary Fig. S3).

**Fig 2:**
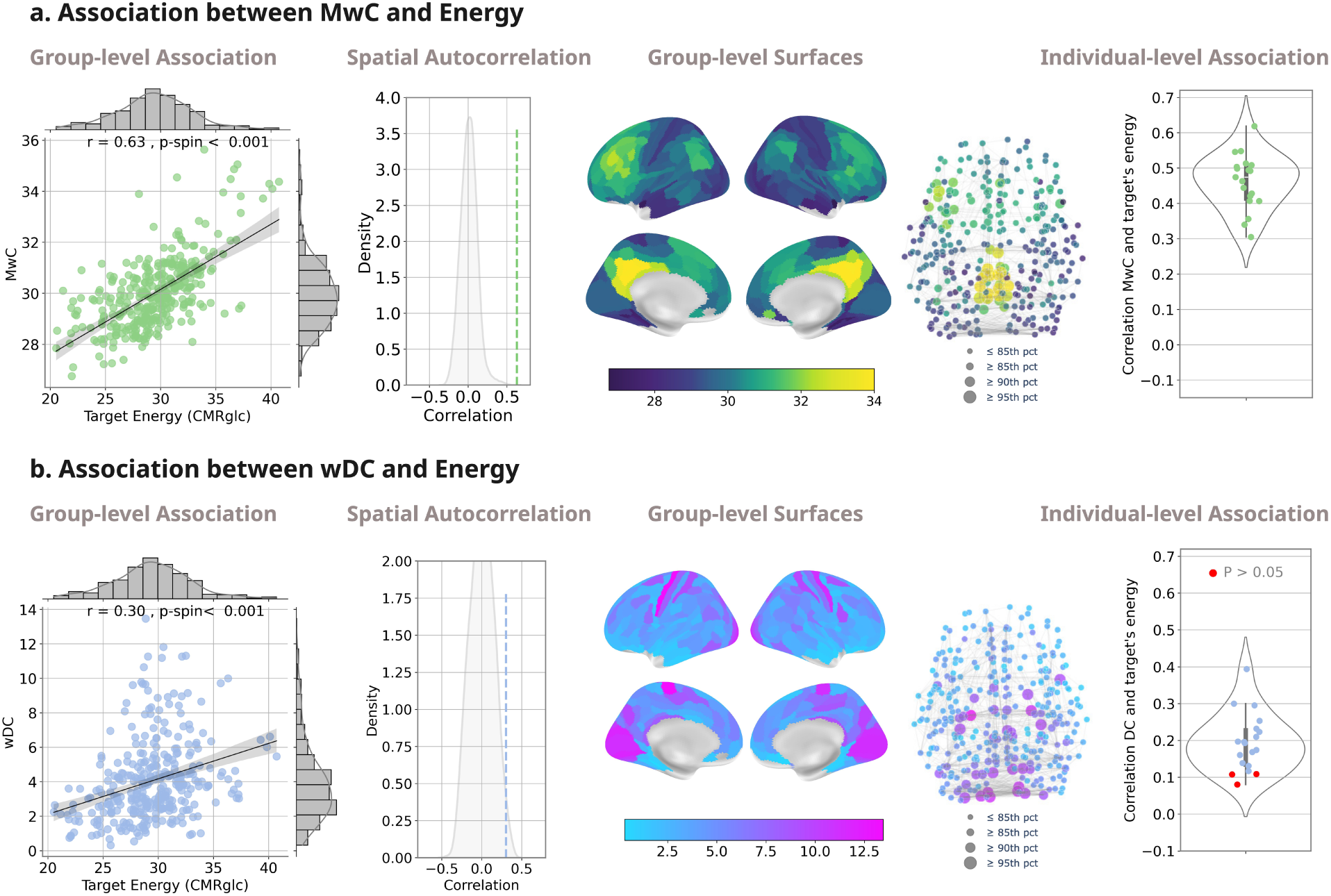
MwC outperforms classical metrics in explaining cortical energy metabolism. **(a)** Association between MwC and CMRglc at the group and individual subject levels. The linear regression plot shows the significant association between calculated MwC and measured CMRglc of target regions across the cortex at the group level. The histogram plot illustrates the null distribution, derived from spin test, while the dashed line represents the empirical correlation, which assesses the influence of spatial autocorrelation. The empirical correlation (dashed line, r = 0.63) lies outside the 90^th^ percentiles (5th–95th percentiles: [-0.22, 0.25]) of the null distribution. The four brain surfaces illustrate the group-level MwC values, with lower values in dark blue and higher values in yellow. The graph plot representation highlights regions with the highest values (≥85^th^, ≥90^th^, and ≥95^th^ percentiles), shown as progressively larger node sizes, and color coding is consistent with the group-level surface maps. The violin plot shows the significant subject-level correlations, where each dot represents the individual correlation between regional MwC and CMRglc. **(b)** Association between wDC and CMRglc at the group and individual subject levels. The linear regression plot reveals a moderate correlation between regional wDC and CMRglc across all brain regions. The histogram plot illustrates the empirical correlation (dashed line, r = 0.31) against a null distribution corrected for spatial autocorrelation (5th–95th percentiles: [-0.16, 0.19]). The brain surfaces illustrate the group-level wDC values across cortical brain regions. The graph plot representation highlights the top 5%, 15%, and 25% of regions with the highest wDC values, represented as increasing node sizes, while maintaining the same color coding as in the group-level surface plots. The violin plot details subject-level correlations, with red dots highlighting nonsignificant correlations.

Additionally, we performed several control analyses. First, we tested for a statistical difference between MwC and wDC in their correlation with CMRglc. Across the joint results from all three datasets (Supplementary Fig. S4), MwC was significantly stronger correlated with CMRglc than wDC (Steiger’s test, Z = 2.54, p = 0.01). A stepwise, multiple linear regression revealed that MwC explained 39.9% of CMRglc variance, while wDC additionally explained 4.2% of variance in energy metabolism. Second, we assessed the relevance of each component in the two-layer network to the sensitivity of the MwC calculation. Removing the E(i) term – thereby reducing MwC to wDC – lowered the group-level correlation from 0.63 to 0.31 (p < 0.001), with individual correlations dropping to 0.10 – 0.27 (all p < 0.05). Removing the effect of nFC for each target by setting identical weights across all connections resulted in a similar reduction in performance (group-level r = 0.33, p < 0.001; individual r = 0.11 – 0.52). Third, we tested whether accurate matching between regional connectivity and metabolism is essential, or if merely scaling wDC with arbitrary metabolism could account for our findings. To assess this, we shuffled the connection weights across the vector of regional energy values. This preserved the regional distribution of energy metabolism and the pattern of brain-wide connectivity but disrupted their true pairing. Across 500 iterations, the correlation values between MwC and CMRglc dropped to around zero at both the group- and subject-level (Supplementary Fig. S5a). This analysis demonstrates that including metabolic data alone is insufficient to explain the observed correlation with CMRglc. Moreover, several sensitivity analyses confirmed that the observed MwC-CMRglc correlations were stable across different thresholding levels and were unaffected by region size (Supplementary Fig. S5b).

Further control analyses confirmed the robustness of the main association between MwC and CMRglc. To evaluate the reliability of the regional parcellation, we repeated the main MwC–CMRglc analysis using the Julich-Brain atlas (28) and obtained similar results (Supplementary Fig. S6). We also tested whether differences in the quality of atlas registration among subjects influenced the subject-level MwC–CMRglc association. Using the correlation ratio (corratio) metric for aligning MNI to functional and PET data, we found no significant associations with the subject-level MwC–CMRglc coupling (MNI-to-functional: r = 0.09, p = 0.72; MNI-to-PET: r = 0.18, p = 0.46) (Supplementary Fig. S7). Next, tested whether parcel-wise variability across subjects in the MNI-registered T1-weighted images influenced the group-level spatial correlation. We computed a voxelwise variance-to-mean ratio (VMR) map (29) across subjects, averaged these values within each parcel, and repeated the MwC–CMRglc analysis after regressing out the resulting parcel-wise VMR map. The main association remained unchanged (raw: r = 0.628; controlled for VMR: r = 0.625). Finally, we repeated the MwC–CMRglc spatial correlation analysis while accounting for the temporal signal-to-noise ratio (tSNR) of the fMRI and PET maps. The main association remained strong after adjusting for regional PET tSNR (r = 0.609) and fMRI tSNR (r = 0.699), indicating that the observed relationship was not influenced by regional differences in signal quality. Having established the physiological validity of MwC at the regional level, we next tested its relationship with the macroscale architecture of cognition.

### Metabolism-weighted hubs align with cognitive network hierarchies

In the following section, we investigated the distribution of MwC across large-scale brain networks and whether MwC indicates the complexity of cognitive domains processed by cortical regions. Fig. 3a shows the group-average regional values of MwC (upper graph) and wDC (lower graph), color-coded according to six major functional brain networks (30) and sorted in ascending order by the networks’ medians. Our findings reveal that MwC significantly increases from sensory (visual, VIS; sensory-motor, SOM) to higher cognitive networks, including the default mode network (DMN), salience/ventral attention (SAL), and control network (CON) (One-way ANOVA, F = 31.36, p-value < 0.001; post-hoc Tukey test: p<0.05). In contrast, the average wDC showed a less distinctive distribution across all networks (F = 2.97, p-value = 0.01). Significant pairwise differences between mean network centrality measures are marked with colored stars, and these were more prevalent in MwC compared to wDC. We also examined the relationship between MwC and wDC with continuous cortical gradients. MwC showed a significant positive relationship with the principal functional gradient (31) (r = 0.485, p < 0.001), whereas wDC displayed a negative association (r = -0.227, p < 0.001). Additionally, MwC was not significantly related to the principal microstructural gradient (32) (r = 0.051, p = 0.36), while wDC showed a significant negative correlation (r = -0.363, p < 0.001) (Supplementary Fig. S8).

**Fig 3:**
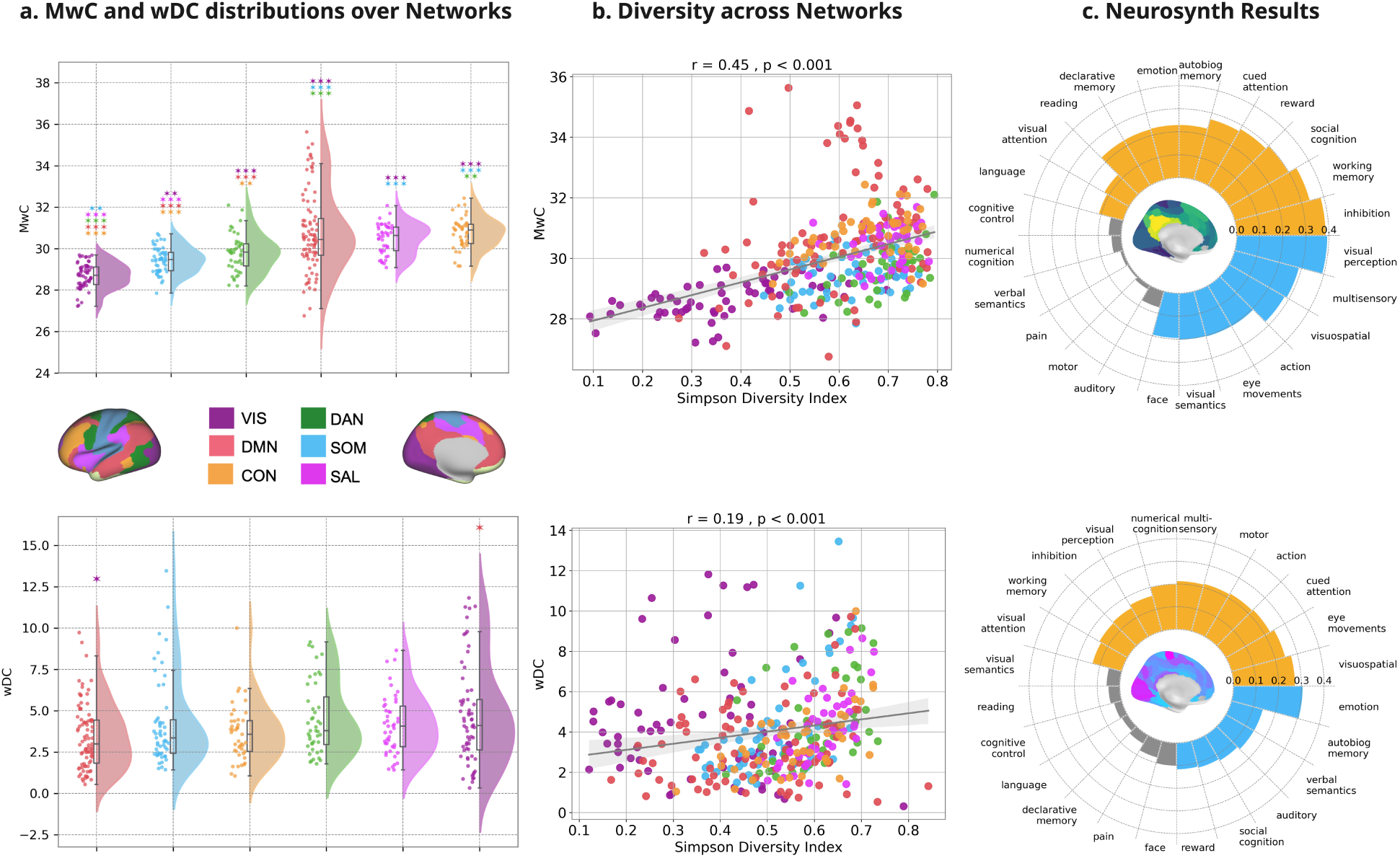
MwC hubs map onto cognitive networks and domains. **(a)** MwC and wDC distribution across macroscale brain networks. Brain regions are sorted by functional brain networks (30). The top row shows the group-level MwC, whereas the bottom row shows the wDC distribution. The brain surfaces illustrate the distribution of six Yeo-networks across the cortex. Both MwC and wDC are ordered in ascending order of median network values. Colored stars indicate significant differences between each pair of networks based on post hoc comparisons (Tukey’s HSD test following one-way ANOVA, * = p<0.05, ** = p<0.01, *** = p<0.001). Different colors of stars correspond to pairwise comparisons of each network with the others, each color representing a specific network. **(b)** Information Diversity of Networks. Simpson’s diversity index indicates the connectivity diversity of brain regions across networks. The upper scatter plot shows a significant positive correlation between MwC and SDI. wDC is less significantly correlated with SDI (lower). **(c)** Cognitive domain mapping. Circular diagrams show correlation values between 24 Neurosynth categories and MwC (upper) and wDC (lower). Orange sectors represent significant positive, and blue sectors negative correlations, while gray sectors indicate nonsignificant correlations. MwC is consistently and significantly correlated with higher cognitive tasks (e.g., inhibition, memory, cognition, attention) and negatively correlated with sensory-motor tasks (e.g., visual, motor). This distinction is less consistent for wDC.

We then evaluated the diversity of regional network connectivity. For each region, we calculated the Simpson Diversity Index (SDI), which indicates whether a region primarily connects to its own network (SDI close to 0) or to multiple networks (SDI close to 1). Our analysis revealed a significant positive correlation between SDI and MwC, and to a lesser extent, with wDC (Fig. 3b). Finally, we compared the regional distribution of MwC and wDC with the distribution of regional activity related to major cognitive functions, as identified in the Neurosynth database (31). We extracted the activity patterns of 24 major cognitive domains and ranked their correlations with MwC (upper section) and wDC (lower section) in the circular diagrams of Figure 3c. Notably, MwC effectively distinguishes between higher cognitive (represented in orange, such as inhibition and memory) and sensory-motor processing (represented in blue). In contrast, wDC did not clearly differentiate between these two categories; for example, tasks related to "visuospatial" and "eye movements" were more associated with regions of higher wDC values rather than lower values. In summary, MwC serves as a robust and distinctive measure of brain network organization and its relationship with cognitive processing, complementing edge-oriented approaches with specific information about signaling hubs within the brain’s connectome.

### Metabolism-weighted hubness reflects synaptic gene expression and susceptibility to neurodegenerative diseases

In the final section, we investigated the biological plausibility of the MwC metric. First, we tested whether independently sampled transcriptomics data confirm the regional levels of signaling activity and metabolism as indicated by the measure of MwC. We mapped gene expression data from the Allen Human Brain Atlas (AHBA) (33) onto the HCP MMP-atlas and used partial least squares (PLS) regression to identify genes significantly expressed in relation to MwC. At the group level, we found a positive correlation (r = 0.51, p-spin = 0.005) between the first PLS component of genes (PLS1) and MwC (Fig. 4a). In addition, ten-fold cross-validation yielded a mean out-of-sample correlation of r = 0.45 ± 0.26. A subsequent gene enrichment analysis of PLS1-genes with positive Z-scores revealed two major gene clusters that were significantly over-represented in regions of high MwC: genes associated with metabolic activity and synaptic signaling (Fig. 4b). Among the metabolic genes, significant clusters were related to ‘mitochondrial membrane‘, ‘transporter complex’ and ‘mitochondrial matrix’. The majority of significant gene clusters related to synaptic signaling included compartments such as ‘presynapse’, ‘postsynapse’, or the ‘dendritic tree’ (see Table 1 in the supplementary for detailed information on each cluster). Control analyses using randomly selected genes confirmed the specificity of these associations (Fig. S9).

**Fig 4:**
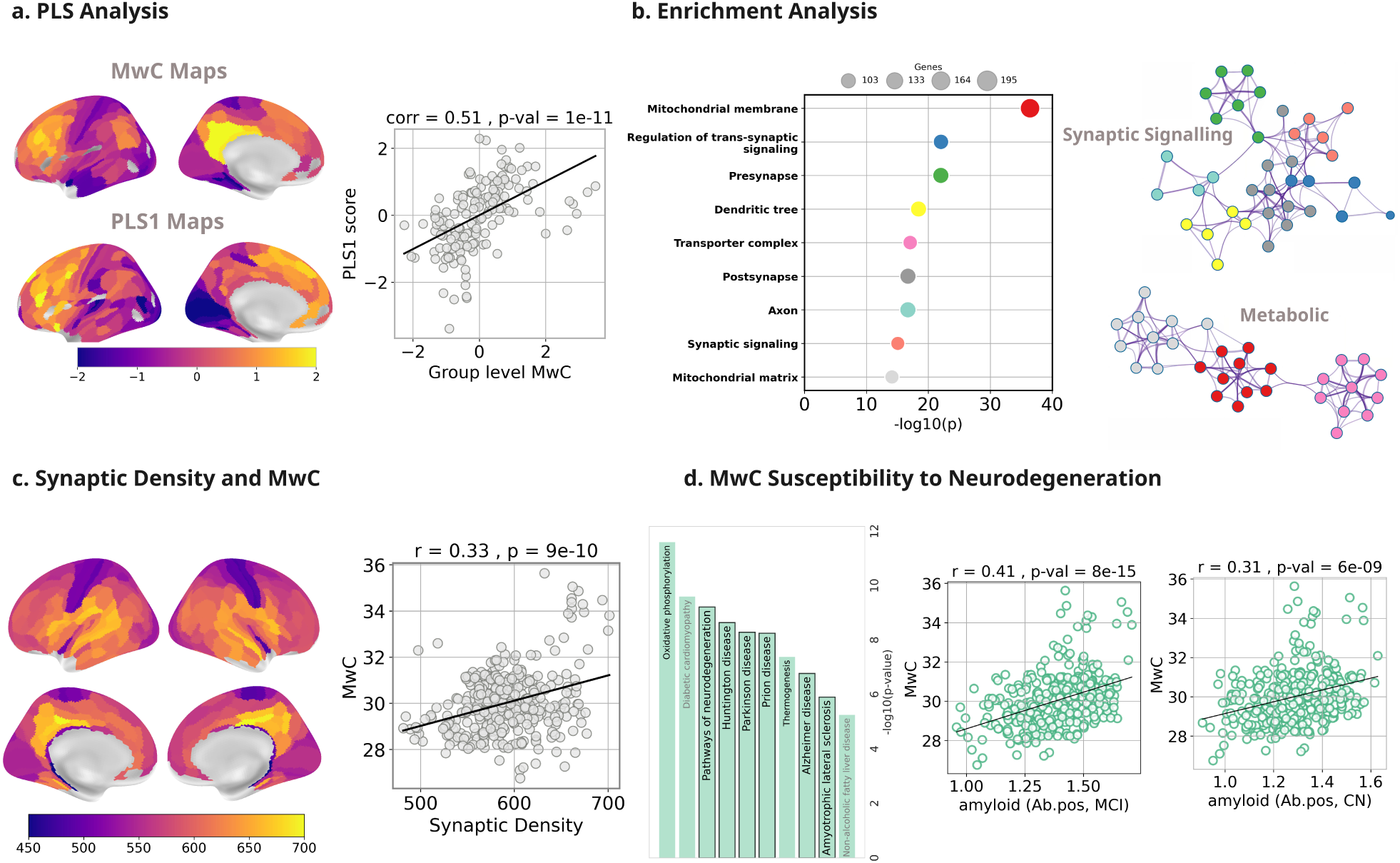
Molecular and pathological correlates of MwC hubness. **(a)** Partial least squares (PLS) analysis was performed to link MwC to gene expression data from the Allen Human Brain Atlas (AHBA), using MwC as the response variable and regional gene expression as regressors. Brain maps display group-level MwC and the first principal component of PLS (PLS1), with a positive correlation between MwC and PLS1 in the scatter plot. **(b)** Gene enrichment analysis for positive Z-score genes using Metascape (73) highlights clusters with shared characteristics, visualized in the left diagram. Clusters with p <1e-10 are shown, with marker size denoting gene counts. The mitochondrial membrane cluster (red) has the most genes and the smallest p-value, emphasizing its significance. The top subnetwork relates to synaptic signaling (e.g., colored clusters for synapses, axons, dendrites), while the bottom predominantly features metabolic clusters**. (c)** Relationship between synaptic density and MwC. Surface maps show regional synaptic density based on SV2A PET imaging ([11C]UCB-J). Warmer colors indicate higher synaptic density. The scatter plot shows a significant positive correlation between synaptic density and MwC across regions (r = 0.33, p = 1e-09), suggesting that regions with higher synaptic density tend to have higher MwC values. **(d)** KEGG pathway enrichment analysis for positive PLS1 Z-score genes identifies the top 10 significant pathways. Brain-related pathways are printed in black, while neurodegenerative disease pathways are emphasized with bold black outlines. Scatter plots illustrate the association between MwC and independent, beta-amyloid imaging data for MCI patients (left) and cognitively normal (CN) subjects (right), revealing a significant positive correlation in each.

Following the transcriptomics analysis results, we explored independent imaging data on synaptic and mitochondrial density in the human brain to evaluate their relationship with MwC. The brain surfaces in Fig. 4c illustrate group-average data of [^11^C]UCB-J PET imaging of the synaptic vesicle glycoprotein 2A (SV2A), which serves as a marker of synaptic density (34). We found a significant positive correlation between synaptic density and MwC across the cortex (r = 0.33, p < 0.001). We also studied the relationship between MwC and mitochondrial density (MitoD) as well as mitochondrial respiratory capacity (MRC) (35), but did not find any significant correlation (see Supplementary Fig. S10).

Finally, we used the KEGG database to identify biological and pathological pathways related to the genes of the PLS1. KEGG analysis revealed “oxidative phosphorylation” as the most significant pathway. Among the nine additional significant pathways identified, six were associated with neurodegenerative diseases (Fig. 4d, bar plot). This suggests that high MwC regions are not only metabolically active but also particularly vulnerable to neurodegeneration. To further validate this finding, we used an external dataset of amyloid-PET imaging, which visualizes amyloid plaques - a hallmark of Alzheimer’s Disease – to determine if high MwC regions are especially prone to the accumulation of amyloid plaques. Consistent with the KEGG analysis results, we found a significant positive correlation between MwC and amyloid load in both cognitively normal (r = 0.32, p < 0.001) and amyloid-β-positive (Ab.pos) MCI patients (r = 0.4, p < 0.001), an early stage of Alzheimer’s disease (Fig. 4d, regression plots). In summary, these findings reveal a cross-scale correspondence: regional energy consumption emerges as the organizing principle that links macroscale connectivity with microscale synaptic machinery. The same energy-intensive hubs that sustain cognitive integration appear most exposed to metabolic stress, bridging network theory and cellular vulnerability.

## Discussion

Network neuroscience has largely characterized the brain as a graph of interconnected nodes and edges, emphasizing the architecture of connections rather than the level of local activity. The present work reframes this view by introducing a metabolism-weighted connectome, an energetically informed model in which both the strength and the metabolic demand of connections determine nodal influence. This framework introduces a physiological dimension to network topology, demonstrating that hubs in the human brain are defined not only by their structural position but also by their activity level. The result is a shift from describing connectivity as a static map to understanding it as a dynamic system constrained by the bioenergetics of signaling.

Critically, MwC explained a larger share of regional glucose metabolism variance than standard centrality metrics, suggesting that the brain’s energy landscape is not merely correlated with network topology, but is constitutively shaped by it. Prior work has shown that cortex-wide FC strength shares organizational properties at molecular and metabolic levels (36), and that connectome hubs are characterized not only by dense anatomical wiring but also by elevated energy metabolism (37). Yet these observations treat metabolism and connectivity as parallel spatial maps that can be compared post hoc. MwC fundamentally extends this framework: rather than aligning a metabolic map onto a connectivity map (19), it embeds regional glucose metabolism directly into the centrality estimation itself. The result is a measure reflecting information throughput rather than connection count alone. MwC hubs therefore identify metabolically dominant regions that sustain integrative processing across their pathways, a concept well-grounded in the principle that the majority of neuronal energy expenditure supports synaptic signaling (38, 20, 39, 22).

MwC also expands upon existing models of brain communication which infer connectivity strength from statistical synchrony of BOLD signals. Since BOLD signal fluctuations reflect relative changes in rather than absolute activity (25, 26), FC can misrepresent regions with high synchrony but low signaling throughput (40–43). By scaling connection strength with metabolic demand, MwC resolves this discrepancy, allowing for a distinction between connectivity that is present and connectivity that is energetically engaged. This biological realism reframes network integration not as a static architectural property, but as an energetically constrained process of communication.

The topology revealed by MwC also carries broader implications for understanding cognitive function. High-MwC regions were concentrated in association networks involved in executive control, attention, and memory, systems that depend on continuous integration of distributed information. By measuring quantitative energy metabolism across the cortex, we and others have identified distinct metabolic profiles in association regions (9, 44). This supports the view that energy allocation is a limiting factor for large-scale cognition. By aligning network theory with metabolic constraints, MwC offers a mechanistic explanation for how the brain balances efficiency and cost: its most integrative regions are also its most energetically taxed.

At the molecular level, transcriptomic and imaging data confirmed that high-MwC hubs show elevated expression of genes related to synaptic signaling and mitochondrial metabolism, and increased synaptic density on SV2A-PET imaging. These convergent findings suggest that regions of high MwC are supported by dense synaptic and metabolic machinery, consistent with their role as signaling nexuses (45–47). However, this same energetic specialization may confer susceptibility to degeneration. The overlap between MwC hubs and regions of amyloid accumulation and mitochondrial gene upregulation (48–51) points to a fundamental trade-off between integrative capacity and cellular resilience. This “metabolic liability” hypothesis aligns with theories that link neurodegenerative vulnerability to lifelong synaptic workload and oxidative stress.

Our study has also several limitations. Despite replication across two independent datasets, the overall N of the three cohorts is modest in size, given the demanding experimental and invasive nature of the imaging setup. This may limit statistical power and the generalizability of effect estimates. A further limitation concerns the gene enrichment analysis, which was performed using Metascape’s predefined random null model and therefore did not preserve the correlation structure and functional dependencies among genes in the empirical gene set. This may increase the risk of false-positive enrichment results.

In summary, the metabolism-weighted connectome redefines hubness as a property of energy-mediated communication rather than structural prominence. By integrating connectivity with local bioenergetic demand, MwC integrates molecular metabolism with systems-level network organization. This biologically grounded perspective offers a framework to investigate how energetic constraints shape cognition, development, and disease. Furthermore, incorporating metabolic parameters into network models could help predict how local energetic bottlenecks limit distributed computation, paving the way for energetically informed theories of brain function.

## Materials and Methods

### Participants

Our analysis incorporates three independent datasets for main results (TUM.main) and replication analyses (TUM.rep, Vienna.rep). All participants were right-handed and did not have any documented psychiatric conditions. Before data collection, participants provided informed consent and were apprised of potential risks. The full study protocol was approved by the ethics committees of Klinikum Rechts der Isar, Technische Universität München, and the Medical University of Vienna. Data acquisitions were conducted with approval from the ethics committees at Klinikum Rechts der Isar, Technische Universität München, and the Medical University of Vienna. The TUM.main dataset included twenty participants, evenly divided between genders, with an average age of 34.45 ± 5 years. The TUM.rep dataset comprised ten subjects with an average age of 27 ± 5, also equally split between genders. The Vienna dataset consisted of five males and five females, with an average age of 27 ± 7 years. For the TUM.main and Vienna datasets, participants kept their eyes open, while participants in the TUM.rep dataset had their eyes closed during resting state data acquisition.

### Data acquisition

Both TUM datasets were acquired on an integrated 3T PET/MR Siemens Biograph scanner with a 12-channel head coil for MRI imaging. PET data were gathered in list-mode format with an average intravenous bolus injection of 184 MBq (s.d. = 12 MBq) of [18F]FDG. Concurrently, every second, automatic arterial blood samples were taken from the radial artery to measure blood radioactivity using a Twilite blood sampler (Swisstrace, Zurich, Switzerland). Functional MRI data were acquired over a 10-minute interval using a single-shot echo planar imaging sequence (300 volumes; 35 slices; repetition time, TR = 2000 ms; echo time, TE = 30 ms; flip angle, FA = 90°; field of view, FOV = 192 × 192 mm2; matrix size = 64 × 64; voxel size = 3 × 3 × 3.6 mm3). Diffusion-weighted images (DWI) were acquired with a single-shot echo planar imaging sequence (60 slices; 30 non-colinear gradient directions; b-value = 800 s/mm2 and one b=0 s/mm2 image; TR = 10800 ms, TE = 82 ms; FA = 90°; FOV = 260 × 264 mm2; matrix size = 130 x 132; voxel size = 2 × 2 × 2 mm3). Anatomical images were based on a T1-weighted 3D-MPRAGE sequence (256 slices; TR = 2300 ms; TE = 2.98 ms; FA = 9°; FOV = 256 × 240 mm2; matrix size = 256 × 240; voxel size = 1 × 1 × 1 mm3). We used the same data as (44); for details on the data acquisition and preprocessing, see this paper.

The data acquisition from the Vienna site followed similar protocols to those used at the TUM site, as previously detailed by (52). Key differences included a higher and more variable dose of injected [18F]FDG (mean dose = 356 MBq, SD = 66 MBq), the manual measurement of blood radioactivity for the arterial input function, and slight adjustments in the fMRI protocol, such as a shorter acquisition time (∼7 min), a repetition time (TR) of 2400 ms, and a voxel size of 2 × 2 × 3.7 mm³.

### MRI processing

The structural and functional MRI data preprocessing was conducted using the Configurable Pipeline for the Analysis of Connectomes (C-PAC, version 1.4.0) following established procedures:

- Anatomical images underwent skull-stripping, segmentation into the cerebrospinal fluid (CSF), white matter (WM), and gray matter (GM), and were registered to the MNI152NLin6ASym template. Individual GM masks were generated based on probability thresholds and temporal signal-to-noise ratios.
- Functional image processing included slice-time correction, realignment, motion correction, skull-stripping, and registration to anatomical images. Global mean intensity was normalized, nuisance signals (scanner drift, physiological noise, head motion) were regressed out, and time series were bandpass filtered.

DWI preprocessing and probabilistic tractography were executed using MRtrix3 (53), FSL, and Advanced Normalization Tools (ANTs), incorporating denoising, eddy-current correction, motion correction (using FSL top-up), and bias-field correction (using ANTs). Structural connectivity (SC) matrices were derived via constrained spherical deconvolution probabilistic tractography. Spherical-deconvolution-informed filtering was applied to pictograms constrained by anatomical tissue masks and the HCP-MMP parcellation. The strength of SC was assessed based on communicability between brain regions, encompassing both direct and indirect connections.

### PET processing

PET imaging data acquisition for both TUM datasets involved offline reconstruction of the initial 45-minute period using the NiftyPET library (54). This reconstruction employed the ordered subsets expectation maximization (OSEM) algorithm with 14 subsets and 4 iterations divided into 33 dynamic frames. These frames consisted of 10 intervals of 12 seconds, 8 intervals of 30 seconds, 8 intervals of 60 seconds, 2 intervals of 180 seconds, and 5 intervals of 300 seconds. Attenuation correction relied on T1-derived pseudo-CT images (55). All reconstructed PET images underwent motion correction and spatial smoothing using a Gaussian filter with a full width at half maximum (FWHM) of 6 mm. The net uptake rate constant (Ki) was determined using the Patlak plot model (56) based on the last 5 frames of the preprocessed PET images (frames occurring between 20 to 45 minutes) and the arterial input function derived from preprocessed arterial blood samples. The cerebral metabolic rate of glucose (CMRglc) was computed by multiplying the K_i_ map with the concentration of glucose in the plasma of each participant, divided by a lumped constant of 0.65 (57). Finally, the CMRglc maps underwent partial volume correction using GM, WM, and CSF masks derived from the T1 images via the iterative Yang method (58). They were registered to the MNI152NLin6ASym 3 mm template through the anatomical image.

### Model description

The MwC model involves two main steps: (1) constructing a fully-weighted graph and (2) calculating the MwC for each brain region.

#### 1. Construction of a fully-weighted graph

All analyses were performed at the regional level using 334 cortical regions from the Human Connectome Project Multi-Modal Parcellation (HCP-MMP) atlas (27). For both modalities (fMRI and PET), the atlas was transformed in two steps: first from MNI space to each subject’s T1-weighted anatomical space, and then from T1 space to the subject’s fMRI or PET space. The resulting subject-specific atlas maps were then used to obtain parcel-wise values in each modality. Regions assigned to the Yeo limbic network were excluded because of susceptibility-related fMRI artifacts. As our model relies on region-wise correspondence between fMRI and CMRglc, the same parcels were also excluded from the CMRglc maps to maintain spatial consistency across modalities.

Each region’s mean BOLD time series was calculated and used to compute pairwise FC via mutual information (MI). MI quantifies the statistical dependence between two regions’ time series *X* and *Y*, capturing both linear and nonlinear relationships. We used Gaussian Copula Mutual Information (GCMI) (59) to estimate MI efficiently, assuming continuous variables transformed to a standard normal distribution. MI was computed as:

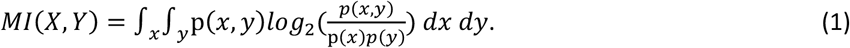

*p(x,y)* is the joint probability density of two arbitrary regions’ time series *X* and *Y*, while *p(x)* and *p(y)* are their marginal densities.

SC was obtained from probabilistic tractography using MRtrix3-based on diffusion-weighted imaging. As our diffusion acquisition was performed within a simultaneous PET-MR protocol rather than as a dedicated high-angular-resolution tractography dataset, we used SC conservatively as a binary anatomical constraint rather than as a quantitative edge-weight measure. Specifically, we retained the top 10% of SC values and binarized the resulting matrix to define the presence or absence of structural connections.

The fully-weighted brain graph was constructed by combining structural, functional, and metabolic information:

- **Edges** were assigned weights equal to the MI between functionally connected regions, but only if a structural connection (from the binarized SC matrix) was present. This was implemented by element-wise multiplication of the SC and MI matrices, resulting FC matrix.
- **Nodes** were annotated with metabolic activity values *E*, defined as the regional median of the CMRglc from FDG-PET data.

This graph forms the basis for calculating the MwC, integrating both regional energy metabolism and the structure-function topology of the connectome.

#### 2. Metabolism-weighted Centrality (MwC)

For each target region *t*, defined as any node for which the metric is computed, the MwC was calculated based on the normalized functional connectivity (nFC) to its connected neighbors and their corresponding metabolic activity.

Let *i ∈ {1,2,…,N}* denote the *N* regions connected to region *t*. The FC between *t* and *i*, denoted *FC(t,i)* was calculated using MI and SC matrices. The metabolic activity of region *i*, *E(i)*, corresponds to its median CMRglc, derived from FDG-PET.

To control for the effect of connection count and total FC strength on MwC, we normalized the FC values for each target region as follows:

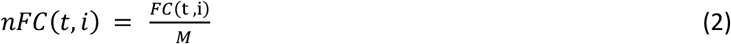

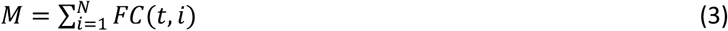

This normalization ensures that:

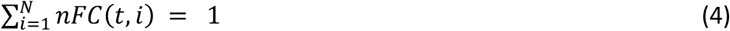

and that the contributions of neighboring nodes are interpreted as proportions rather than absolute values. This prevents target regions with more or stronger raw FC values from yielding artificially inflated MwC scores. Using these normalized weights, MwC was defined as the weighted sum of the metabolic activity of the connected neighbors:

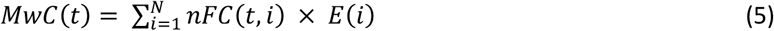

By this definition, MwC is high when a target region is connected to neighbors that have both high metabolic activity *E(i)* and high normalized FC values *nFC(t,i)*. Conversely, MwC is low when neighbors have low metabolic activity and low nFC weights. The CMRglc of the target region itself is excluded from the calculation, avoiding direct dependence between the predictor and outcome in correlation analyses.

### Weighted Degree Centrality (wDC)

wDC was calculated using a conventional edge-weighted graph based solely on FC. First, we found the Pearson correlation between mean BOLD time series of region pairs, consistent with prior studies that have commonly used this approach. The resulting matrix was proportionally thresholded at 20% and then masked using a binarized SC matrix (top 10%) to retain only anatomically plausible connections. We call the final matrix as the FC matrix. For each target region *t*, wDC was computed as:

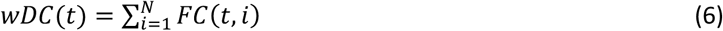

where *N* is the number of connected neighbors and *FC(t,i)* is the connection strength between nodes *t* and *i*.

### Participation Coefficient (PC)

PC was calculated to quantify the extent to which each region distributed its connections across different modules. We used the same structurally constrained and thresholded FC matrix as in wDC analysis. Community structure was identified using the Infomap algorithm (60), as previously applied in brain networks (61, 62). Then, the Infomap algorithm was applied to the FC matrix for each subject. For a given target region *t*, PC was defined as:

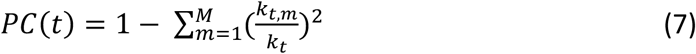

where *M* is the total number of detected modules, *k_t,m_* is the sum of FC weights between region *t* and all regions in module *m*, and *k_t_* is the total degree of region *t*.

Then regional PC values were averaged across subjects for group-level analysis. Their association with regional CMRglc was evaluated using spatial autocorrelation-corrected permutation testing.

### Control analyses

To evaluate the contribution of each model component, we systematically deleted the effect of each component from the MwC definition and generated a surrogate MwC, then we found the relation between the MwC_sur_ and CMRglc.

First, we removed the metabolic term *E(i)*, reducing the model to:

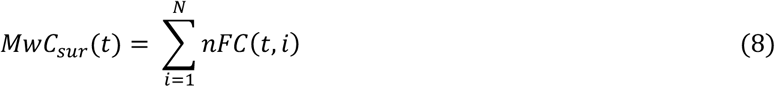

which is equal to one based on equation 4 for all nodes and is therefore uninformative. To create meaningful surrogate data, we instead used raw FC values, which makes MwC similar to the wDC definition in equation 6, with one main difference: here, we have an MI-based FC matrix.

Then, we generated surrogate data by excluding the contribution of nFC data and preserving the same *E*(*i*) values as in the actual data. We manipulated the *nFC* values by assigning equal values to all connections, such that each connection’s strength is set to 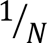 if a target has *N* connected regions, since the sum of all connected regions’ nFC values is equal to one (equation 4):

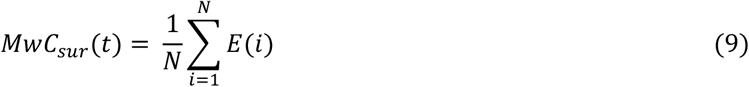

This version corresponds to the average CMRglc of connected neighbors and removes the influence of varying FC strength.

For both surrogate metrics, we computed their correlation with regional CMRglc to assess the effect of removing or flattening each component.

To test whether the observed MwC–CMRglc association depends on the specific alignment between FC weights and neighboring metabolic values, we performed a permutation-based analysis. For each region, we preserved the set of connected neighbors and their CMRglc values but randomly reassigned the FC weights across them. Specifically, for each region *t*, the values *FC(t,i)* were randomly shuffled across the set *i=1,…,N*, MwC was then recomputed as:

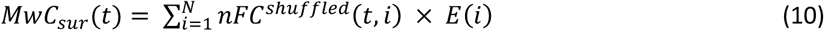

This procedure was repeated 500 times per subject. For each iteration, we computed the correlation between MwC_sur_ and true CMRglc. The resulting distribution served as a null model (Fig. S5a).

We also conducted sensitivity analyses by altering specific model variables and monitoring their effect on the main result, the correlation between MwC and the target’s energy. We analyzed three variables: the SC threshold, the MI threshold, and the target region size. We explored different threshold levels for the first two variables to assess their effect on the correlation between MwC and CMRglc at the group level. We defined the target size as the number of voxels within each target region. To control for regional differences in data quality, we computed regional tSNR maps for both fMRI and PET. fMRI tSNR was calculated from the BOLD time series, whereas PET tSNR was calculated in PET space from the last five frames used for CMRglc estimation. In both modalities, tSNR was defined as the voxelwise mean signal divided by its temporal standard deviation and then averaged within each atlas parcel. We then reassessed the MwC–CMRglc spatial correlation after regressing out these regional tSNR measures.

### Statistical analyses

All statistical analyses were conducted at the regional level. We used Pearson correlation to assess the association between each metric and the regional cerebral metabolic rate of glucose (CMRglc), both at the group level (after averaging across subjects) and at the individual level.

To evaluate the statistical significance of spatial correlations while accounting for spatial autocorrelation, we used spin permutation test (63) which generates null brain maps that preserve the spatial autocorrelation structure of the empirical data. For each test, we generated 10.000 spatially autocorrelated surrogate maps of CMRglc and calculated their correlation with the metric of interest. A two-sided p-value was computed based on the rank of the observed correlation within the null distribution.

To assess differences across networks in Fig. 3a, we first conducted a one-way ANOVA to test for overall significant differences. Then, to see the pairwise differences among networks, the post hoc Tukey’s HSD (Honestly Significant Difference) test compared the means of each network.

To determine whether the relationship with CMRglc significantly differed between MwC and wDC, we applied Steiger’s Z-test for dependent correlations (64). This test compares two correlation coefficients that share a common variable (CMRglc) within the same dataset. To ensure adequate statistical power for this test, we combined the z-scored values from our three datasets, resulting in a total sample size of 45. Finally, we evaluated the unique and shared contributions of MwC and wDC in explaining variability in CMRglc using linear regression analysis. We implemented two models: one that included only MwC as a predictor (CMRglc ∼ MwC) and another that included MwC and wDC as predictors (CMRglc ∼ MwC + wDC). This approach allowed us to quantify the additional variance explained by wDC beyond what is captured by MwC alone.

### Simpson Diversity Index (SDI)

SDI, also known as the Gini-Simpson Index, quantifies diversity by measuring the probability that two entities randomly selected from a dataset belong to the same type (65). Mathematically, it is defined as:

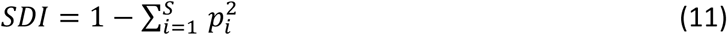

In this context, *S* represents the total number of networks, which amounts to 6 distinct networks. The symbol *p_i_* denotes the proportional abundance of each network *i*. We individually calculated the diversity of *SDI* for each target region. The goal was to quantify the diversity of networks within the connected regions. For a specific target region, *p_i_* indicates the number of connected regions that belong to network type *i*, relative to the total number of connections. The *SDI* index ranges from 0 to 1: a value of 0 indicates no diversity (meaning all entities are of a single type), while a value of 1 denotes maximal diversity, where each network type is equally represented.

### Neurosynth analysis

Neurosynth is a platform that automates the large-scale synthesis of fMRI data (66). It utilizes a database of thousands of published fMRI studies to generate meta-analytic maps based on frequently recurring key terms. This innovative tool enables an objective and quantitative exploration of brain activity associated with various cognitive and sensory processes. Our analysis, particularly as shown in Fig. 3, utilized brain maps for 24 cognitive topic terms derived from previous studies (31, 67). These topics range from lower sensory and motor functions to higher cognitive functions. We examined the correlation between each of these 24 maps and the group-level MwC and wDC maps to explore the cognitive implications of each metric.

### Gene expression analysis using AHBA data

We analyzed the regional expression of 15,631 genes. Regional microarray expression data were obtained from 6 post-mortem brains (1 female, ages 24.0–57.0, 42.50 ± 13.38) provided by the Allen Human Brain Atlas (AHBA, https://human.brain-map.org; (33)). Data were processed with the abagen toolbox (version 0.1.3; https://github.com/rmarkello/abagen) using a 360-region volumetric atlas in MNI space. First, microarray probes were reannotated using data provided by (68); probes not matched to a valid Entrez ID were discarded. Next, probes were filtered based on their expression intensity relative to background noise (69), such that probes with intensity less than the background in ≥50% of samples across donors were discarded, yielding 31,569 probes. When multiple probes indexed the expression of the same gene, we selected and used the probe with the most consistent pattern of regional variation across donors, i.e., differential stability (70). The MNI coordinates of tissue samples were updated to those generated via non-linear registration using the Advanced Normalization Tools (ANTs; https://github.com/chrisfilo/alleninf). Samples were assigned to brain regions in the provided atlas if their MNI coordinates were within 2 mm of a given parcel. To reduce the potential for misassignment, sample-to-region matching was constrained by hemisphere and gross structural divisions (i.e., cortex, subcortex/brainstem, and cerebellum) (68). All tissue samples not assigned to a brain region in the provided atlas were discarded. Inter-subject variation was addressed by normalizing tissue sample expression values across genes using a robust sigmoid function (71). Normalized expression values were then rescaled to the unit interval. Gene expression values were then normalized across tissue samples using an identical procedure. Samples assigned to the same brain region were averaged separately for each donor and then across donors, yielding a regional expression matrix with 360 rows, corresponding to brain regions, and 15,631 columns, corresponding to the retained genes.

#### Partial Least Squares (PLS)

We used partial least squares (PLS) regression to examine the relationship between the group-level MwC map and regional gene expression across the left hemisphere. The predictor matrix consisted of AHBA-derived expression values for 15,631 genes across 180 cortical regions, and the response variable was the corresponding 180 × 1 group-level MwC vector. Before analysis, both the gene-expression matrix and the MwC map were z-scored. PLS was implemented to identify latent components maximizing covariance between regional gene expression and MwC, and analyses focused on the first component (PLS1), which captured the dominant transcriptional pattern associated with MwC.

To assess statistical significance while accounting for spatial autocorrelation, we used a spin-test–based null model with 10,000 iterations. Specifically, the cortical MwC map was rotated on the spherical surface, the same parcel exclusions were then applied, and the PLS analysis was repeated for each rotated map to generate a null distribution of explained variance. The significance of the observed PLS effect was assessed relative to this null distribution.

To estimate the contribution of individual genes to PLS1, we performed 1,000 bootstrap resamples of regions with replacement. For each resample, PLS was rerun and gene weights were aligned to the sign of the original solution. We then computed a bootstrap Z score for each gene as the ratio of its original weight to the standard deviation of its bootstrap weights, and genes were ranked according to these Z scores for downstream enrichment analysis (72).

As an additional robustness analysis, we performed 10-fold cross-validation. In each fold, a one-component PLS model was trained on 90% of cortical regions and evaluated on the remaining 10%, and this procedure was repeated across all folds. Out-of-sample performance was quantified as the correlation between predicted and observed MwC values in the held-out regions.

#### Gene enrichment analysis

We used the Metascape (http://metascape.org) (73), an automated meta-analysis tool, for our gene enrichment analysis. We input the genes exhibiting positive z-score values from the first PLS component into the Metascape website. We tested whether the input gene list shared enrichment pathways with GO “Molecular Functions”, “Biological Processes”, and “Cellular Components” pathways. We used the default parameters on the Metascape website. Metascape’s input limit is 3000 genes; we selected the top 3,000 genes with positive Z score values for our analysis. Within the Metascape results illustration, pathways are ordered based on their p-value. We included the connected clusters and removed the small, isolated clusters.

We also did the enrichment analysis for the KEGG 2021 Human pathway separately using Enrichr (74–76) to find the relationship of our MwC map with disease pathways. We used the same gene list as the Metascape.

### Synaptic density atlas

We used the high-resolution in vivo human brain synaptic density atlas provided by the Neurology Research Unit (NRU) at the University of Copenhagen (34). This atlas was generated using PET imaging with [^11^C]UCB-J, a radioligand that binds to synaptic vesicle glycoprotein 2A (SV2A) and provides quantitative estimates of synaptic density. Data were collected from 33 healthy adult participants (17 females, 16 males), aged 20-70 years (44.8 ± 14.6 years). The atlas represented the group-average synaptic density distribution across subjects. The in vivo PET data were calibrated to absolute concentration values (pmol/mL) using in vitro autoradiography from postmortem human brain tissue. Further details are described in (34).

### Mitochondrial maps

We used mitochondrial phenotype maps from (35) that quantify regional mitochondrial density and respiratory capacity across the human brain. These maps were derived from a coronal slab of the right hemisphere of a 54-year-old neurotypical male, which was physically voxelized into 703 cubes (3 × 3 × 3 mm). Biochemical assays measured mitochondrial content (citrate synthase and mtDNA, combined into MitoD) and oxidative phosphorylation (OXPHOS) enzyme activities (averaged into TRC), from which a tissue-specific mitochondrial respiratory capacity map (MRC = TRC/MitoD) was computed. The maps were initially created in the native anatomical space of the donor brain and subsequently linearly registered to MNI ICBM152 space for neuroimaging compatibility. In our analysis, we resampled our MMP atlas to this MNI space and used it to parcellate the mitochondrial maps.

### Amyloid-beta maps

We used amyloid PET data from the Alzheimer’s Disease Neuroimaging Initiative (ADNI), provided by the authors of a recent study by (77). PET scans were acquired using the AV45 (Florbetapir) tracer, registered to T1-weighted MRIs, and parcellated with the MMP1 atlas using a CAT12-based surface reconstruction pipeline. SUVR values were normalized to the whole cerebellum and extracted for cortical ROIs. To avoid longitudinal confounds, only the first available PET session per subject was included. The final dataset comprised 129 cognitively normal (CN) individuals (mean age = 75.5 ± 7.6 years; 83 females) and 95 individuals with mild cognitive impairment (MCI) (mean age = 76.0 ± 7.4 years; 41 females). These regional SUVR values were used to assess the spatial association between amyloid accumulation and MwC.

## Data availability

All raw and preprocessed data from the TUM site are available for download in the OpenNeuro repository (https://openneuro.org/datasets/ds004513). All further derivative maps for analysis, including the Vienna site data, the β-amyloid maps as described in (77), and the 24 Neurosynth maps, are available on our GitHub repository (https://github.com/NeuroenergeticsLab/metabolic_weighted_centrality). Human gene expression data can be accessed via the Allen Human Brain Atlas database (http://human.brain-map.org/static/download). Synaptic density maps are described in (34) and can be obtained from https://xtra.nru.dk/SV2A-atlas/.

## Code availability

All scripts, Python Jupyter notebooks, and configuration files used to perform the analyses are available on GitHub (https://github.com/NeuroenergeticsLab/metabolic_weighted_centrality). Mutual information calculation was done using GCMI package (https://github.com/robince/gcmi). The code for PLS analysis is available at https://github.com/SarahMorgan/Morphometric_Similarity_SZ. Gene enrichment analyses were performed using Metascape (https://metascape.org). KEGG analysis was conducted with Enrichr (https://maayanlab.cloud/Enrichr/).

## Acknowledgements

We would like to thank Roman Belenya, Dr. Samira Epp, Dr. Antonia Bose, and Dr. André Hechler for their valuable input during the development of this idea. We thank Prof. Thomas Beyer and Lalith Kumar Shiyam Sundar for sharing the Vienna dataset and Dr. Nicolai Franzmeier for sharing the beta-amyloid maps.

## Author contributions

MA designed and implemented the model, performed all analyses, and wrote the manuscript. LF provided guidance on transcriptomics and gene enrichment analyses, their biological interpretations, and revised the manuscript. GC provided supervision, guidance in data analysis and visualization, and revised the manuscript. VR conceptualized the study, contributed to model development, supervised the project, and wrote the manuscript.

