## Supplementary Figures and Tables for "Metabolism-weighted brain connectome reveals synaptic integration and vulnerability to neurodegeneration"

By Ashrafi et al.

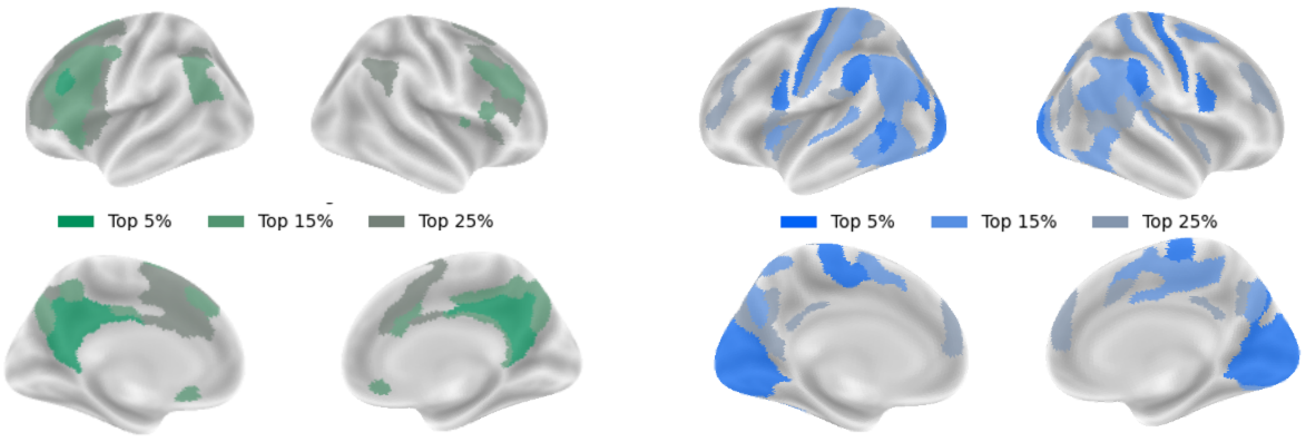


| cortex | Network | Region | Lobe |
|---|---|---|---|
| PCC | Default | RSC_L | Par |
| PCC | Default | 7m_L | Par |
| PCC | Default | POS1_L | Par |
| PCC | Default | v23ab_L | Par |
| PCC | Default | d23ab_L | Par |
| PCC | Default | 31pv_L | Par |
| DLPFC | Cont | p9-46v_L | Fr |
| PCC | Default | 31pd_L | Par |
| PCC | Default | 31a_L | Par |
| PCC | Default | 7m_R | Par |
| PCC | Default | POS1_R | Par |
| PCC | Default | 23d_R | Par |
| PCC | Default | v23ab_R | Par |
| PCC | Default | d23ab_R | Par |
| PCC | Default | 31pv_R | Par |
| PCC | Default | 31pd_R | Par |
| PCC | Default | 31a_R | Par |

| cortex | Network | Region | Lobe |
|---|---|---|---|
| Primary_Visual | Vis | V1_L | Occ |
| Early_Visual | Vis | V2_L | Occ |
| Early_Visual | Vis | V3_L | Occ |
| SomSen_Mot | SomMot | 4_L | Fr |
| PCC | Default | POS2_L | Occ |
| Dors_Str_Vis | Vis | IPS1_L | Par |
| PCL and MCC | SomMot | 24dd_L | Fr |
| SomSen_Mot | SomMot | 2_L | Par |
| Primary_Visual | Vis | V1_R | Occ |
| Early_Visual | Vis | V2_R | Occ |
| Early_Visual | Vis | V3_R | Occ |
| SomSen_Mot | SomMot | 4_R | Fr |
| PCC | Cont | POS2_R | Par |
| Dors_Str_Vis | DorsAttn | IPS1_R | Par |
| PCL and MCC | SalVentAttn | 23c_R | Par |
| SomSen_Mot | SomMot | 2_R | Par |
| PCC | Vis | DVT_R | Par |

**Fig S1**: Comparative analysis of hub regions derived from MwC and DC. The brain surfaces represent the hub regions across two neuroimaging-derived maps to highlight the differences between these maps. Proportional thresholding, including the top 5%, 15%, and 25%, was applied to each map to identify hubs, with x% indicating the most prominent x% values. The tables below the maps show the details of each measure's top 5% hubs. The MwC hub map (left maps), varying in threshold levels, uncovers hubs primarily in regions associated with higher-order cognitive functions in the parietal and frontal lobes. The 5% hubs are in the PCC cortex and DLPFC. The DC hub maps (right ones) identify hubs concentrated in the occipital and some parts of the parietal lobes. Most regions are VIS regions, and those located in the parietal lobe are the SOM regions. It also contains some PCC regions.

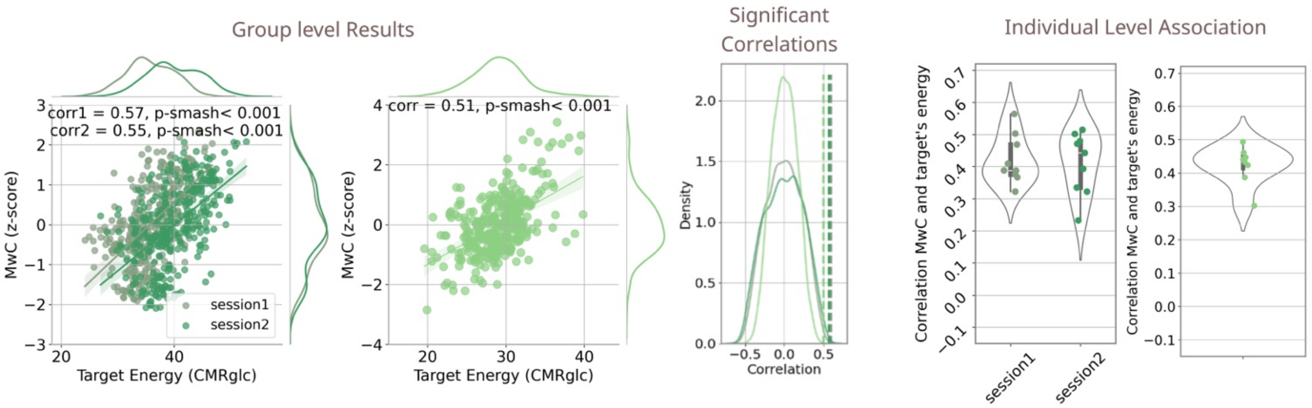


**Fig S2:** The replicability of the MwC model was evaluated using two FDG-PET/MRI datasets: two sessions of the Vienna dataset and the replicated TUM dataset (TUM.rep). Scatter plots illustrate significant group-level correlations between MwC and regional energy for both datasets (r=0.57, p-smash < 0.001 in session 1; r=0.55, p-smash <0.001 in session 2; r = 0.51, p-smash< 0.001 in TUM.rep). The empirical correlations (dashed lines) fall outside of the 90th percentile of the null distribution histograms, shown in the distribution plot. Violin plots show individual-level correlations, ranging from 0.32-0.56 in session 1, 0.23-0.51 in session 2, 0.30- 0.49 in TUM.rep (all p <0.05).

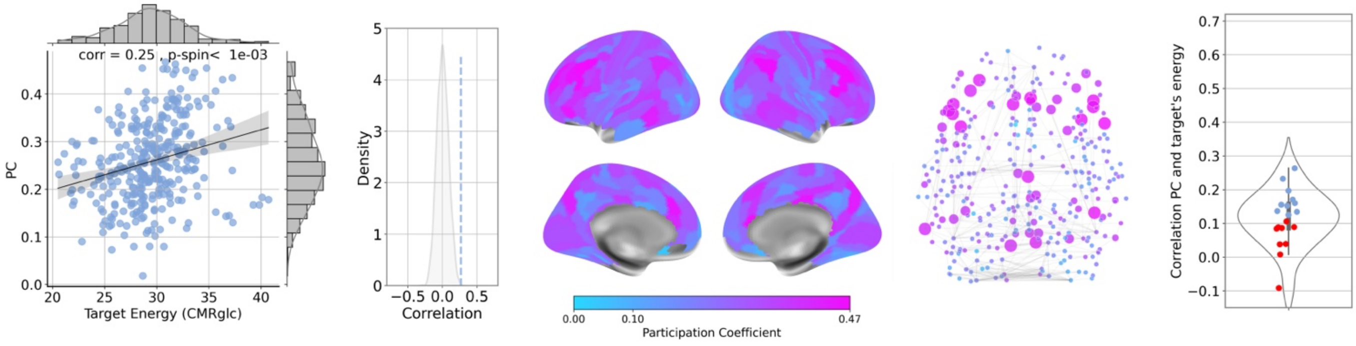


**Figure S3:** Replication of Fig. 2 analyses for participation coefficient (PC) metric.

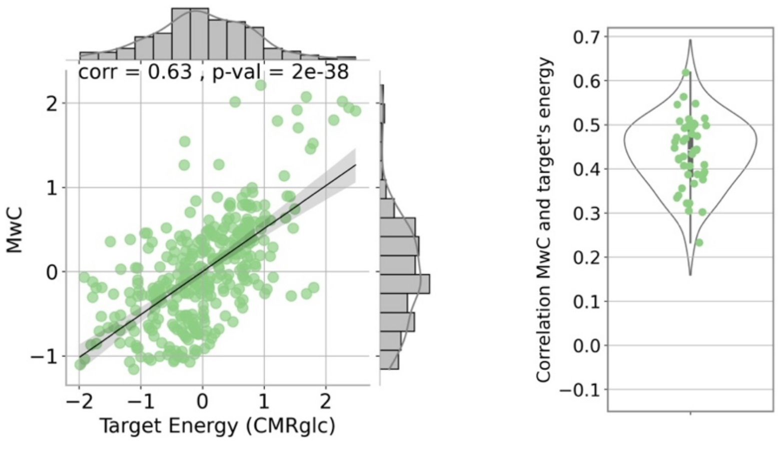


**Figure S4.** Association between MwC and regional metabolic activity at the group and individual subject levels, using the combined data. We concatenated the zscore of three datasets and found the group level MwC across all datasets and subjects. The linear regression plot shows the significant association between MwC and the actual energy of targets across the cortex at the group level. The violin plot shows the significant subject-level correlations, where each dot represents the individual correlation between MwC and regional energy

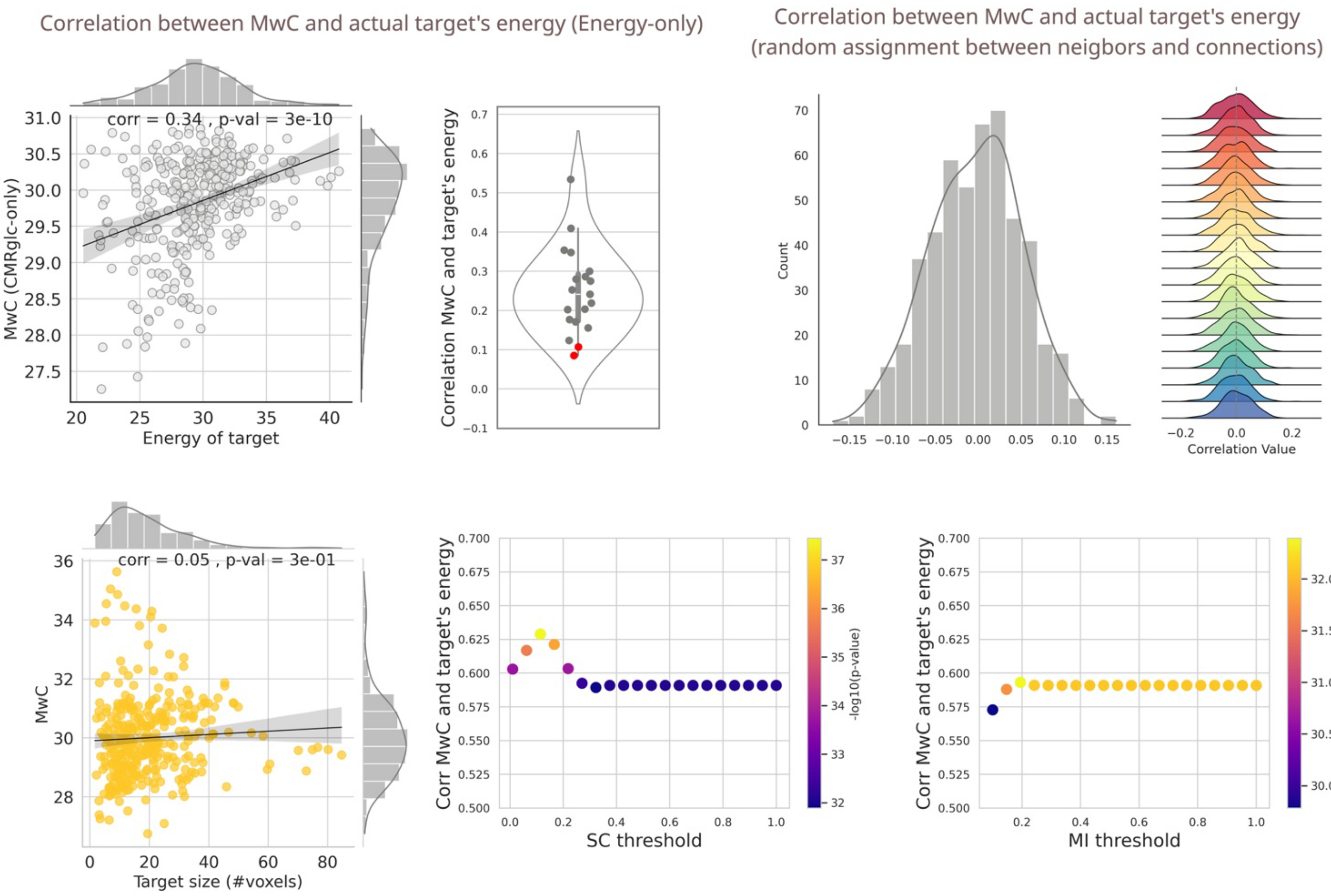


**Figure S5. (a)** Control analyses assessed the contribution of functional connectivity (nFC(t,i)) data to MwC performance. Two plots on the left represent the effects of removing nFC from the MwC definition, resulting in a correlation of 0.33 ($p<0.0001$) at the group level and a reduction in individual-level correlations, with two non-significant correlations shown in red. Two plots on the right show the effect of pairing between E and nFC on the MwC performance, which was tested using random assignments of connection weights to energy values (500 iterations). The histogram illustrates the distribution of group-level correlation values across 500 iterations, centered at zero. The joy plot shows individual-level correlations also centered around zero, with each distribution corresponding to a specific subject. **(b)** Sensitivity analysis. **Left:** Target regions exhibit varying sizes in terms of voxel count. We assessed whether the size of the target region affects the MwC value. Using group data, correlating MwC with target size revealed a non-significant correlation, indicating that MwC is independent of the number of voxels within each target region. **Middle:** We used SC to mask connections and define MwC for each target region. Before defining the SC mask, we applied proportional thresholding to the SC matrix, changing the threshold level from 0.05 to 1, which indicates considering 5% of the biggest connections to 100% of the connections. This figure demonstrates the effect of varying threshold levels of the SC matrix on the correlation between MwC and CMRglc. This thresholding changes the correlation value for threshold levels below 0.3, while the correlation remains consistent beyond this level. **Right:** We employed MI to measure the connection weights between different brain regions. Proportional thresholding was applied to assess the impact of different MI threshold levels on the correlation between MwC and CMRglc. Our result reveals that the correlation between MwC and CMRglc is robust to the MI threshold level.

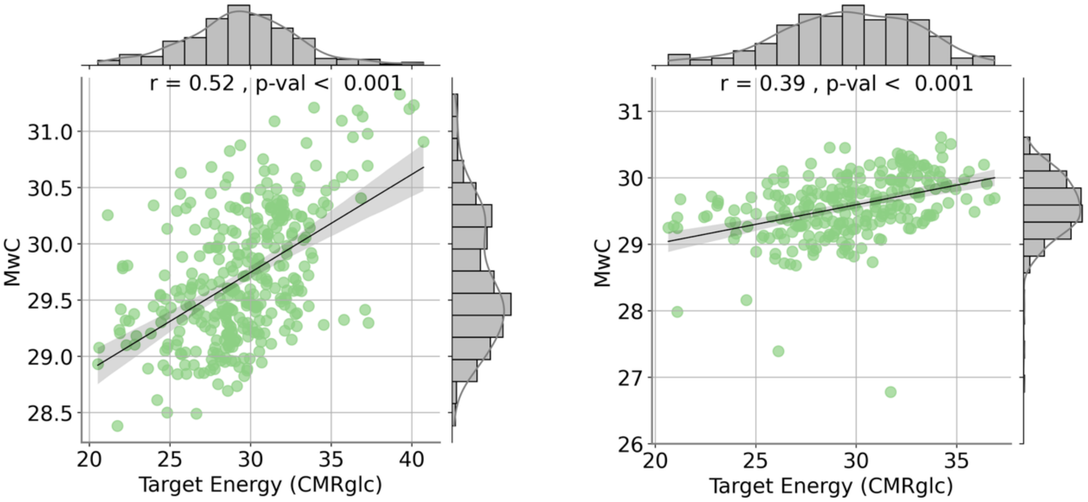


**Figure S6.** *Group-level association between MwC and CMRglc. MwC was calculated without applying the SC binary mask, using the MMP atlas (left) and the Julich Brain atlas (right). Because SC matrices were generated only for the HCP-MMP pipeline, this comparison was performed without SC masking in either atlas.*

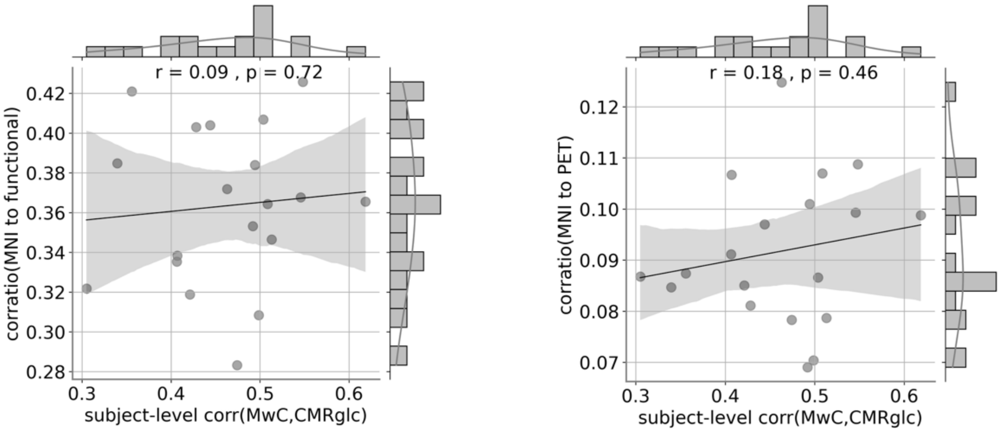


***Figure S7. Registration quality does not explain subject-level MwC–CMRglc coupling.*** *Scatterplots show the association between the subject-level correlation of MwC and CMRglc and registration quality, quantified using the* ***corratio*** *metric, for MNI-to-functional (left) and MNI-to-PET (right) alignment. Neither association was significant.*

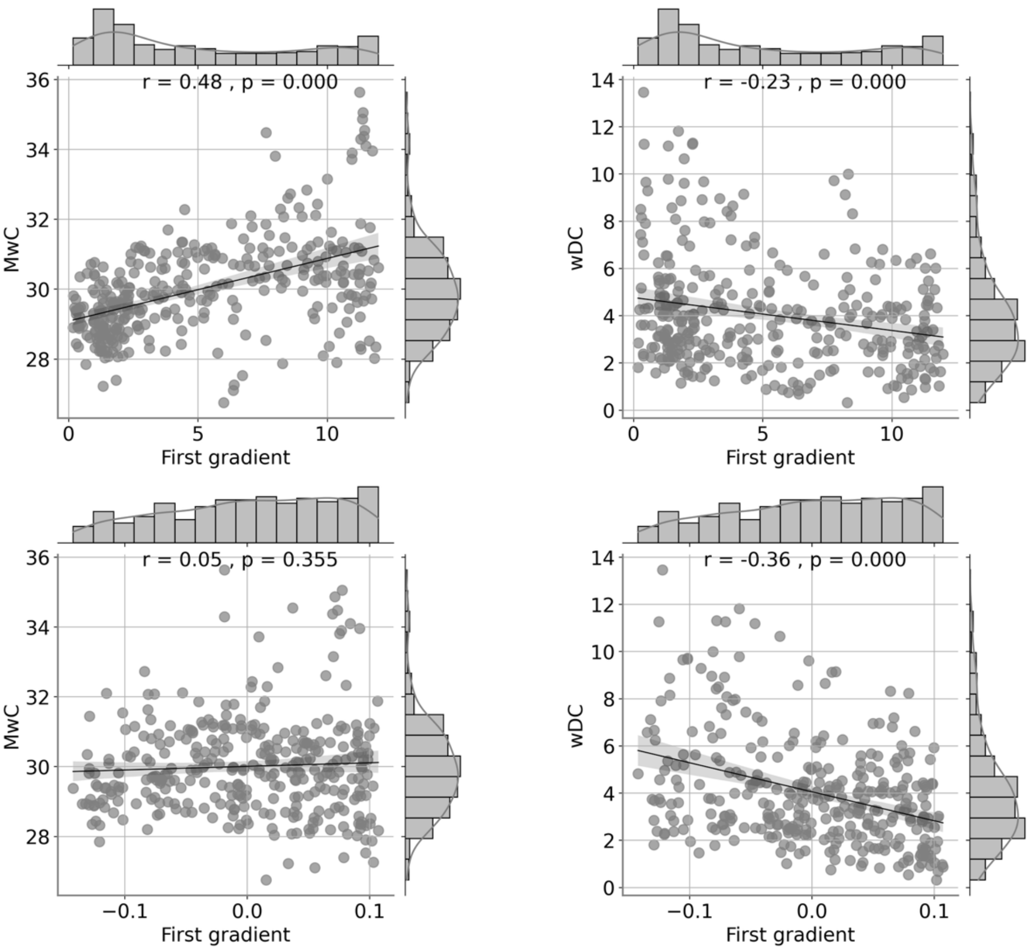


***Figure S8.*** *Relationships of MwC and wDC to functional and microstructural cortical gradients. Scatterplots show region-wise correlations of MwC and wDC with the principal functional gradient (top) and the principal microstructural gradient (bottom). MwC was positively associated with the functional gradient but not with the microstructural gradient, whereas wDC showed negative associations with both gradients.*

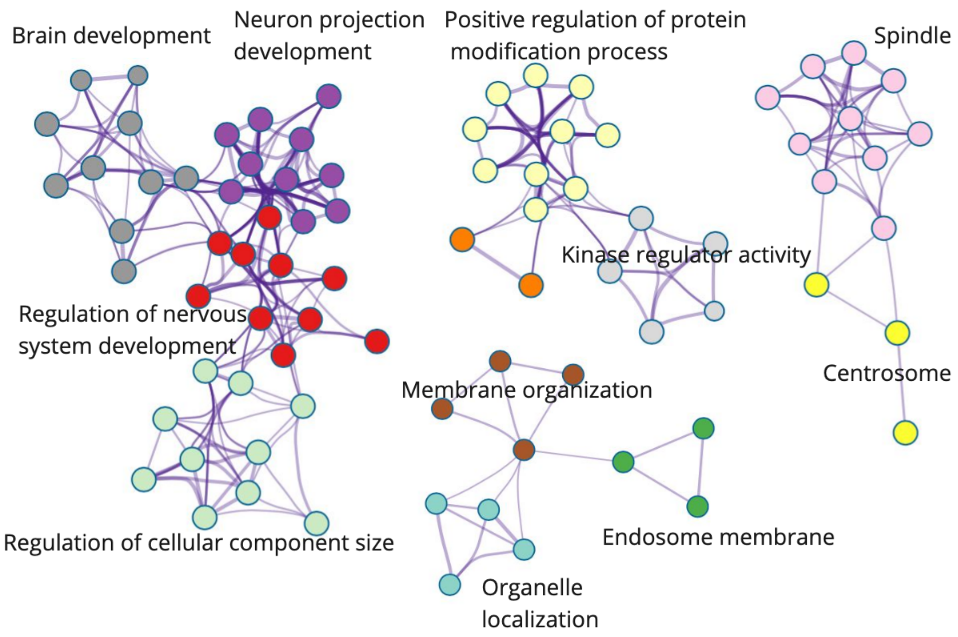


**Figure S9.** Metascape network of random gene set. We selected a random set of genes and imported this set on the Metascape website and subsequently performed enrichment analysis. The categories substantially vary from the PLS analysis shown in Fig. 4, particularly showing more diverse clusters than those on energy metabolism and synaptic signaling.

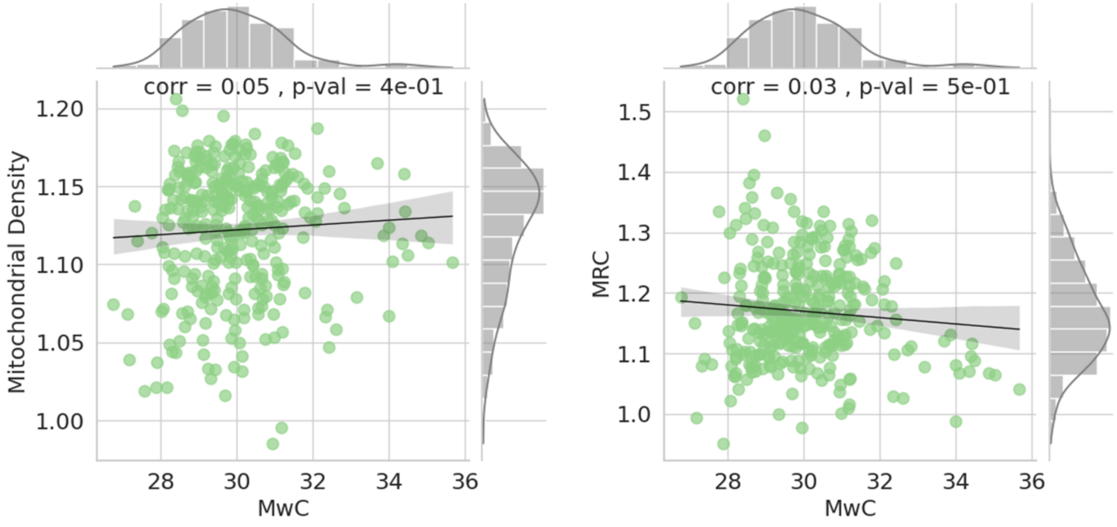


**Figure S10.** The association between group-level MwC with MitoD and MRC. We found no significant association between MwC with MitoD and MRC maps.

**Table 1.** Details of the gene enrichment analysis corresponding to Figure 4b and c. Each row represents a specific pathway from Figure 4b, listing the genes included in that pathway. For the mitochondrial membrane pathway, only genes with a p-value smaller than -10 are shown due to the high number of associated genes.

| | Description | Log(p-value) |
|---|---|---|
| **1. mitochondrial membrane** | **mitochondrial membrane** | -32,07 |
| | organelle inner membrane | -29,69 |
| | mitochondrial inner membrane | -29,30 |
| | mitochondrial protein-containing complex | -25,46 |
| | Pathways of neurodegeneration - multiple diseases | -18,79 |
| | inner mitochondrial membrane protein complex | -18,69 |
| | mitochondrion organization | -18,32 |
| | Oxidative phosphorylation | -16,76 |
| | Huntington disease | -15,91 |
| | Diabetic cardiomyopathy | -15,28 |
| | Parkinson disease | -14,91 |
| | generation of precursor metabolites and energy | -14,91 |
| | Prion disease | -14,91 |
| | Alzheimer disease | -14,39 |
| | Thermogenesis | -12,85 |
| | Amyotrophic lateral sclerosis | -12,25 |
| | energy derivation by oxidation of organic compounds | -12,19 |
| | carbohydrate derivative biosynthetic process | -11,50 |
| | Chemical carcinogenesis - reactive oxygen species | -11,08 |
| | cellular respiration | -11,07 |
| | aerobic respiration | -11,00 |
| | nucleoside phosphate metabolic process | -10,92 |
| | nucleobase-containing small molecule metabolic process | -10,74 |
| | nucleotide metabolic process | -10,66 |
| | mitochondrial respiratory chain complex assembly | -10,64 |
| | oxidative phosphorylation | -10,54 |
| | purine nucleotide metabolic process | -10,27 |
| | purine-containing compound metabolic process | -10,18 |
| **2. regulation of trans-synaptic signaling** | regulation of trans-synaptic signaling | -22,04 |
| | modulation of chemical synaptic transmission | -21,62 |
| | regulation of synaptic plasticity | -11,21 |
| | positive regulation of synaptic transmission | -5,18 |
| | long-term synaptic potentiation | -2,50 |
| **3. presynapse** | presynapse | -22,02 |
| | transport vesicle | -13,38 |
| | exocytic vesicle | -9,66 |
| | transport vesicle membrane | -7,59 |
| | synaptic vesicle | -7,37 |
| | synaptic vesicle membrane | -6,28 |
| | exocytic vesicle membrane | -6,28 |
| **4. dendritic tree** | dendritic tree | -18,42 |
| | dendrite | -18,12 |
| | cell body | -16,41 |
| | neuronal cell body | -12,62 |

| | | |
|---|---|---|
| | perikaryon | -8,54 |
| | transporter complex | -17,07 |
| | transmembrane transporter complex | -16,25 |
| | monoatomic ion transmembrane transporter activity | -9,50 |
| | monoatomic cation transport | -9,48 |
| | monoatomic ion channel activity | -8,46 |
| | plasma membrane protein complex | -8,30 |
| | monoatomic ion channel complex | -8,18 |
| | inorganic ion transmembrane transport | -7,90 |
| | channel activity | -7,70 |
| | monoatomic cation transmembrane transport | -7,70 |
| | passive transmembrane transporter activity | -7,66 |
| | monoatomic cation transmembrane transporter activity | -7,43 |
| | inorganic molecular entity transmembrane transporter | -7,38 |
| | activity | -7,15 |
| | inorganic cation transmembrane transport | -7,10 |
| | gated channel activity | -6,80 |
| | inorganic cation transmembrane transporter activity | -6,79 |
| **5. Transporter complex** | cation channel complex | -6,46 |
| | monoatomic cation channel activity | -6,10 |
| | potassium channel complex | -6,09 |
| | metal ion transport | -5,49 |
| | voltage-gated potassium channel complex | -5,02 |
| | voltage-gated monoatomic ion channel activity | -4,97 |
| | voltage-gated channel activity | -4,80 |
| | salt transmembrane transporter activity | -4,32 |
| | voltage-gated monoatomic cation channel activity | -3,94 |
| | potassium channel activity | -3,16 |
| | potassium ion transmembrane transporter activity | -2,87 |
| | voltage-gated potassium channel activity | -2,55 |
| | potassium ion transmembrane transport | -2,42 |
| | potassium ion transport | |
| | postsynapse | -16,71 |
| | glutamatergic synapse | -13,73 |
| | neuron to neuron synapse | -11,75 |
| | asymmetric synapse | -11,19 |
| | postsynaptic specialization | -10,83 |
| | postsynaptic density | -10,51 |
| | synaptic membrane | -8,72 |
| | presynaptic membrane | -6,2 |
| | postsynaptic membrane | -5,49 |
| | dendritic spine | -5,08 |
| **6. postsynapse** | neuron spine | -4,97 |
| | postsynaptic specialization membrane | -3,65 |
| | postsynaptic density membrane | -3,28 |
| | axon | -16,7 |
| **7. axon** | distal axon | -10,14 |
| | growth cone | -5,52 |
| | site of polarized growth | -5,21 |

| | | |
|---|---|---|
| **8. Synaptic signaling** | synaptic signaling | 15,07 |
| | trans-synaptic signaling | 14,38 |
| | chemical synaptic transmission | 14,21 |
| | anterograde trans-synaptic signaling | 14,21 |
| | Neuroactive ligand-receptor interaction | -5,71 |
| **9. Mitochondrial matrix** | mitochondrial matrix | -14,13 |
| | amide biosynthetic process | -10,56 |
| | peptide metabolic process | -9,62 |
| | peptide biosynthetic process | -8,54 |
| | organellar ribosome | -7,24 |
| | mitochondrial ribosome | -7,24 |
| | translation | -7,16 |
| | mitochondrial translation | -6,91 |
| | organellar large ribosomal subunit | -6,11 |
| | mitochondrial large ribosomal subunit | -6,11 |
| | ribosomal subunit | -5,80 |
| | mitochondrial gene expression | -5,23 |
| | structural constituent of ribosome | -5,2 |
| | ribosome | -4,66 |
| | large ribosomal subunit | -4,20 |
| | Ribosome | -2,47 |
